# Automation of Complex Patient-Derived Organoid (PDO) Culture and Screen

**DOI:** 10.64898/2026.01.02.697417

**Authors:** Hannah A Strobel, Jonathan C Hoying, Sarah M Moss, Victoria Marsh Durban, Maria R Oyaga, Angeline Lim, Wienand Omta, Victor Wong, David A Egan, Hozefa Amijee, James B Hoying

**Affiliations:** Advanced Solutions Life Sciences, Manchester NH, USA; Cellesce Ltd., Cardiff, UK; Core Life Analytics, ‘s-Hertogenbosch, NL; Molecular Devices, San Jose, CA, USA

**Author notes:** corresponding author: James B. Hoying.

**Keywords:** organoid, spheroid, automation, high-throughput, drug screening, patient-derived organoid, image-based profiling

## Abstract

Tissue organoids and spheroids are powerful tools for many different applications, particularly in the area of drug screening. Yet, their widespread use is limited by a need for automated cell and tissue culture procedures and increased throughput. Here, we demonstrate that the BioAssembyBot® (BAB) automation platform, with integrated BioStorageBot® (BSB) and imaging system can perform tasks normally done by scientists with increased efficiency and reduced error. Using the BAB Hand®| Pipette tool, BAB can precisely seed a Matrigel® organoid suspension such that Matrigel® domes are formed consistently in the center of each well plate. The BAB-automated process performed faster, with a higher success rate, and less organoid fragmentation than when the process is done by an experienced scientist. We then repeated this workflow with patient-derived colorectal cancer organoids (CRC PDOs) and performed a fully automated drug screen on cultured organoids. Image-based profiling performed on images that were automatically acquired was able to distinguish clear differences in patient response to anticancer drugs between two donors. Overall, we demonstrate the capability of the BioAssemblyBot® automation platform to manage an entire organoid drug screening protocol with minimal human interaction. The system is user-friendly and may be adapted to a wide variety of workflows and applications, providing a solution to the need for organoid automation technology.

## Introduction

Organoids and spheroids, including patient-derived models, are promising, useful tools in drug target discovery and screening. The ability to scale up organoid production, culture, and assaying are important in their widespread utility. Automation is key, not only for increasing throughput, but also for improving consistency and reducing the risk of human error during fabrication and use. We have previously demonstrated that with the robotic BioAssemblyBot Platform, adherent cells can be automatically passaged, cell suspensions can be dispensed into multiple well plates, and advanced 3D tissue models can be automatically fabricated, cultured, and assayed^1^. This platform is comprised of the 6-axis robotic BioAssemblyBot® (BAB), the BioStorageBot modular incubator, and the intuitive BioApps™ workflow control software, which is integrated with confocal imaging. Here, we use the BAB Platform to fully automate a tumor organoid/spheroid-related workflow using either HCT-116-derived spheroids or in a patient-derived colorectal cancer organoid (PDO) drug screen assay. The workflows involved seeding domes of Matrigel organoids/spheroids into wells of a 96-well plate, incubating in an integrated, modular incubator (BioStorageBot®); performing daily culture medium changes; treating with drug dilutions; and performing staining and automated imaging procedures.

Initial studies compared HCT-116 colorectal cancer spheroid morphology in cultures managed by the automated system to those managed by expert, standard manual practices. Then, a drug screen was performed on patient-derived colorectal cancer organoids. PDOs are being increasingly used for both cancer research and drug testing because they better represent the heterogenous patient biology than more traditional cell lines^2, 3^. PDOs have enormous potential for use in customizing medications for each individual patient^4-6^. A multitude of anti-cancer drugs can be tested on each patients’ individual tumor cells to predict which course of treatment will be most effective for that particular patient.

To evaluate drug effects on PDOs, we measured organoid viability using the ATP CellTiter-Glo® assay. However, this approach only provides overall viability readout at the well level and lacks information on the PDO morphology. To further detect changes in PDOs in response to drug exposure, we applied image-based cell profiling methods^7^ to quantify phenotypic changes in the PDOs in an unbiased manner. Using high-content imaging, measurements related to PDO size, shape, and texture were obtained for each organoid. Thus, further enriching the information available at the organoid level.

Here, we applied our automation workflow to CRC PDOs, performing a fully automated drug screen on PDOs treated with the anti-cancer drugs Trametinib and Adavosertib. Following the automated preparation, feeding, drug treatment, and image acquisition, image sets were analyzed to generate phenotypic responses of the PDO to the drugs. All measurements from the image data were used to generate morphological profiles which were then compared to the control (untreated PDOs) profiles. We were able to demonstrate that not only is the BAB automation platform faster and more efficient than human scientists, but also that the automated PDO assay with image-based profiling was able to discriminate differences between PDOs treated with the different drugs more clearly than traditional statistical methods.

## Methods

In this study, we evaluated the feasibility of using the BioAssembly Platform to plate and culture PDOs in a fully automated workflow. First, we used HCT spheroids in a proof-of-concept study comparing the effects of automated culture on organoid plating efficiency and growth.

Then, we obtained CRC PDOs from two separate donors and performed a drug screen to determine which chemotherapeutic was most effective for each donor. This was assessed by measuring both organoid viability and performing an image-based profiling protocol with deep learning.

### The BioAssemblyBot Automation Platform

The BioAssembly Platform (**Figure 1**) consists of Advanced Solutions Life Sciences (ASLS) BioAssemblyBot®400 (BAB400), containing a 6-axis robotic arm capable of utilizing a wide range of interchangeable bioprinting, fluid dispensing, material movement, and tissue fabrication tools. Additionally, our modular BioStorageBot incubator and the ImageXpress® Confocal HT.ai High-Content Imaging System from Molecular Devices were integrated to complete the manufacturing platform for this application. For this specific spheroid workflow, the platform employed a BioAssemblyBot BAB Hand™ | Pipette, a BAB Hand™ | Pick & Place, and modular tube, pipette tip, cooling, and pipetting stations. All these components and the operational tasks to complete the workflow, including automated communication between the different instruments, are controlled by our user-friendly BioApps software, a platform that connects, controls, and instructs the platform to perform each step in a workflow. Steps are shown in **Figure 2**.

**Figure 1:**
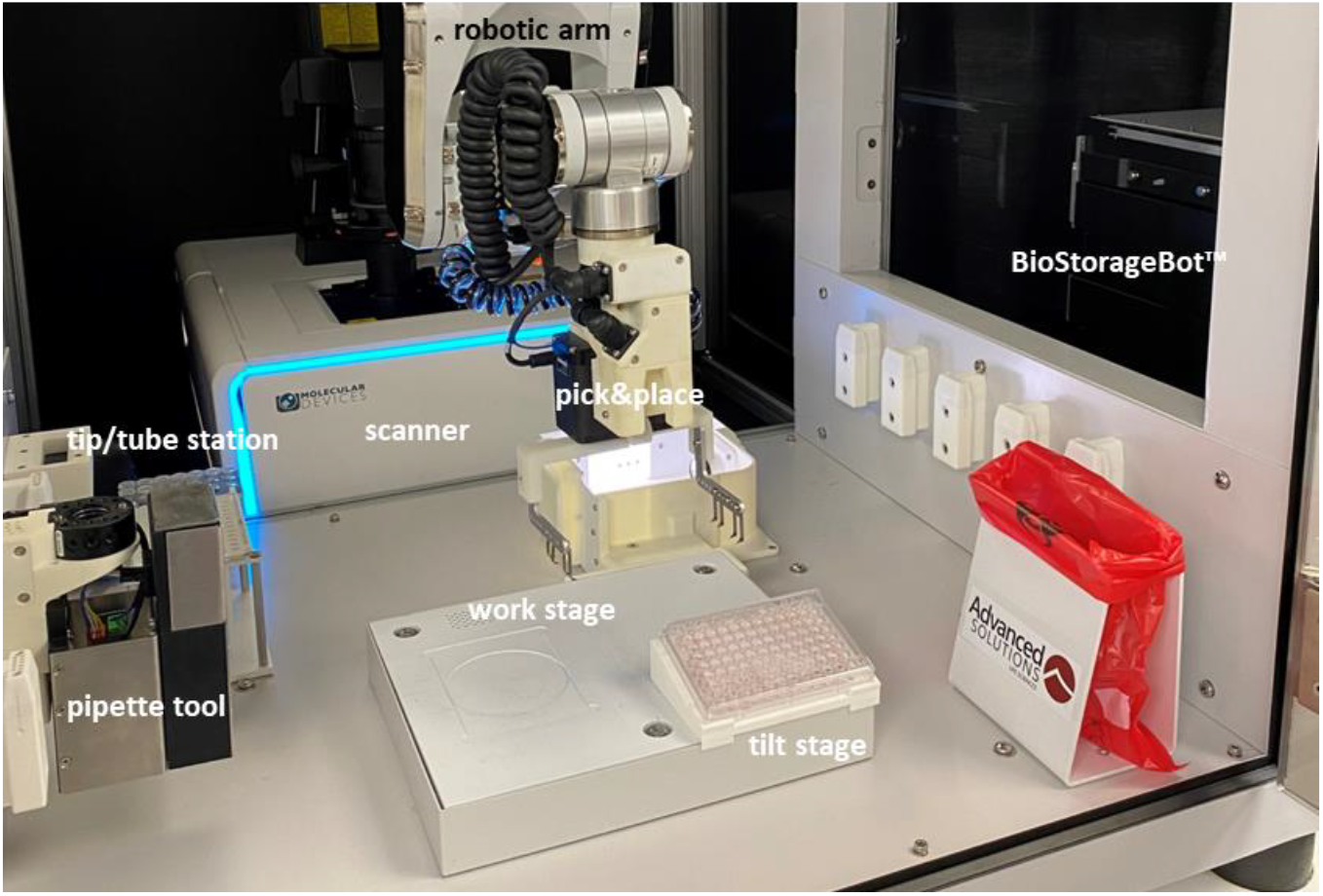
BioAssemblyBot Platform with integrated high content imager (ImageXpress confocal HT.ai system) and a modular incubator (BioStorageBot).

**Figure 2:**
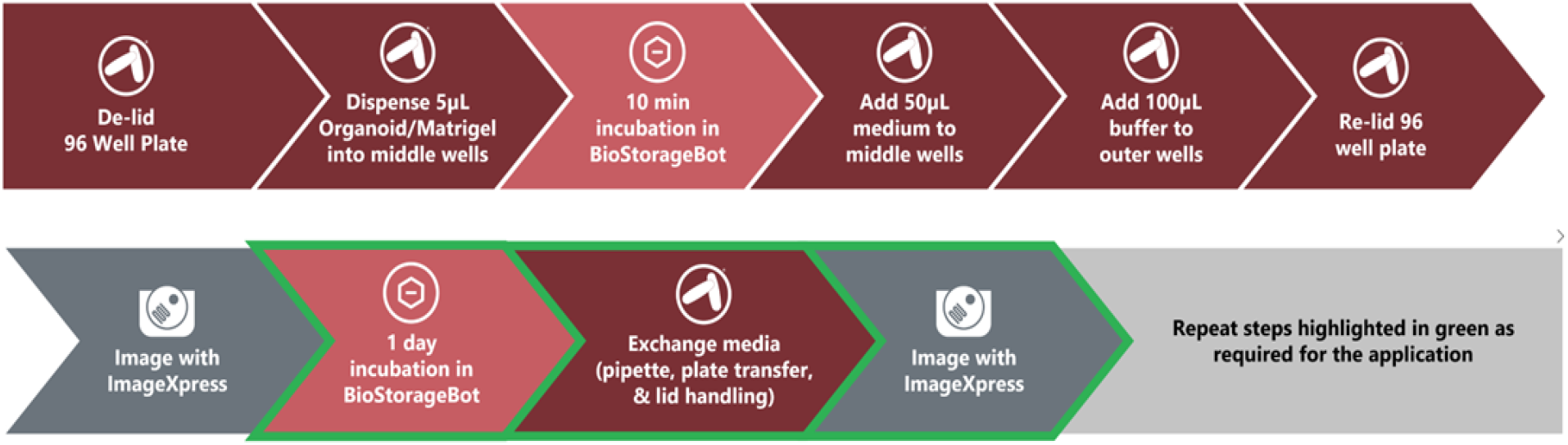
Schematic of the spheroid/organoid workflow automated with the BioAssembly Platform.

### HCT-116 Organoid Seeding and Culture

For preliminary experiments examining organoid handling and consistency, tumor spheroids were pre-formed manually by plating HCT-116 colorectal cancer cells (ATCC cat# CCL-247) into a non-adherent v-bottom plate (Greiner) at a concentration of 2,000 cells per well. After 24 hours, spheroids were suspended in 80% Matrigel in culture medium (EMEM, Gibco, with 10% FBS). 96 spheroids per 250 µl of cold Matrigel were suspended in a sterile tube and placed within the cooling station within the platform. Using the BAB Hand™ | Pipette, the BAB400 aspirated from the chilled tube and then dispensed 5 µl of the spheroid/Matrigel suspension into the center of wells of a 96-well plate to form discernable domes on the bottom of each well. Only the central wells of the plate were used to avoid known edge effects of the outer wells (the outer wells were filled with buffer).

After dispensing, a 10-minute wait period allowed the Matrigel domes to gel before 50 µl of culture medium was added to each well, followed by transfer to the BioStorageBot for culturing. Each day, as managed by the tumor-spheroid BioApp, the BioAssembly Platform automatically performed culture medium exchanges and acquired transmitted light images daily via the ImageXpress Confocal HT.ai system (details below). For medium exchanges, the plate was transferred to a modular tilt stage within the platform that holds the well plate at an angle, analogous to someone holding the plate at a tilt when manually pipetting. This allows for more complete culture medium removal from each well without disturbing the Matrigel dome in the center.

### PDO Seeding and Culture

3D Ready® (Molecular Devices) CRC PDOs were obtained from two different donors. PDOs were suspended in 80% Matrigel and dispensed into a 96-well plate using the BAB and Pipette Hand. 5µl were dispensed into the center of each well as described above, such that a “dome” is formed. After gelling, BAB added culture medium to each well and moved the plate to the modular BioStorageBot incubator. On Day 2 of culture, BAB changed culture media and added a serial dilution series of the MEK1/2 inhibitor Trametinib (Selleckchem) or the WEE1 inhibitor Adavosertib (Selleckchem) to wells. Final Trametinib concentrations were 25, 10, 2.5, or 1 nM, and final Adavosertib concentrations were 5µM or 50, 5, or 0.5nM. On Day 7, cultures were imaged using an integrated confocal imager (Incell) following a Hoechst staining protocol. Hoechst dye (Life Technologies) was diluted 1:3000 in diH_2_O and incubated on cultures for 6 minutes at RT before 3 rinsing steps with PBS. All steps of the PDO assay, including staining and automated imaging protocol, were controlled by a user-defined PDO BioApp.

### CellTiter-Glo 3D Assay

After imaging, a CellTiter-Glo 3D assay (Promega) was performed on plates to measure ATP production. Briefly, the CellTiter-Glo 3D reagent was added to plates containing Matrigel-embedded organoids, which were incubated for 30 min at RT on a shaker plate. Supernatants were then removed to a white 96-well plate and read using a luminescence filter on a BioTek® plate reader.

### Image Acquisition for PDO Study

Image acquisition of PDOs was carried out on the INcell confocal scanner. PDOs were stained with Hoechst dye (Invitrogen) at RT. After 6 min of incubation, the staining solution was replaced with media and organoids were imaged live (in brightfield and DAPI). To capture the 3D PDO samples, confocal imaging was carried out using the 10X objective, with Z-stacking enabled (Z-step size 4µm). Maximum intensity projection images were saved and used for downstream image analysis.

### Image Analysis and Data Analytics

Image analysis was performed using IN Carta® Image Analysis Software (Molecular Devices). Briefly, each PDO in the image was segmented in the brightfield channel using a custom deep learning-based model trained in the IN Carta software (ver1.17). A decision tree classifier was applied to the analyzed data as a quality control step (e.g. organoids with saturated pixels were excluded). Features for size, texture, and channel intensities were extracted from all channels (brightfield, FITC and DAPI). A total of 76 measurements for each PDO were then exported for data analysis.

StratoMineR™ software (Core Life Analytics, Utrecht NL), a cloud-based analytics platform was used for hit selection and cluster analysis. Briefly, data was uploaded to the StratoMineR platform and the following processing steps were applied: Data transformation, feature scaling, dimensionality reduction, hit selection, and cluster analysis^8^. Principal component analysis (PCA) was used to reduce the data to three components followed by hit selection which is based on the calculated Euclidean distance score for the components.

## Results

### Workflow Efficiency

Successful dome seeding is defined as the formation of discernable domes that did not spread out into a thin disc on the well bottom. From this, 81.25% of wells contained discernable domes when dispensed by BAB400, compared to 72.0% when manually performed by an experienced scientist. The entire workflow for seeding domes, which includes dispensing domes, gelation (10 min wait), dispensing media to domes, dispensing buffer to empty outer wells, and transfer to the BioStorageBot® incubator, took 26 minutes and 46 seconds for the BioAssembly system, compared to 41 minutes and 24 seconds when performed manually. The daily tasks of moving the plate of domes to and from the BioStorageBot for media exchanges and moving to and from the ImageXpress Confocal HT.ai system took 13 min and 40 sec (not including scan time).

### HCT Organoid Morphology

From the daily images (**Figure 3A**), the average size of individual HCT spheroids was determined to be approximately half the diameter when seeded manually compared to those plated by BAB400 (**Figure 3B**). It was also noted that an average of 6.2 ± 2.8 spheroids were present in each dome formed manually in contrast to 2.0 ± 1.2 spheroids per dome formed by BAB400, the expected number based on seeding density. Combined, this suggests that the spheroids were fragmented by manual operations, likely due to inconsistent pipetting rates when performed by the scientist, but not by the BAB Platform. Over time, spheroids in both groups steadily increased their overall diameter (**Figure 3B**). Calculation of spheroid growth rates indicated that individual spheroids increased in size at similar rates for both workflows (**Figure 3C**).

**Figure 3:**
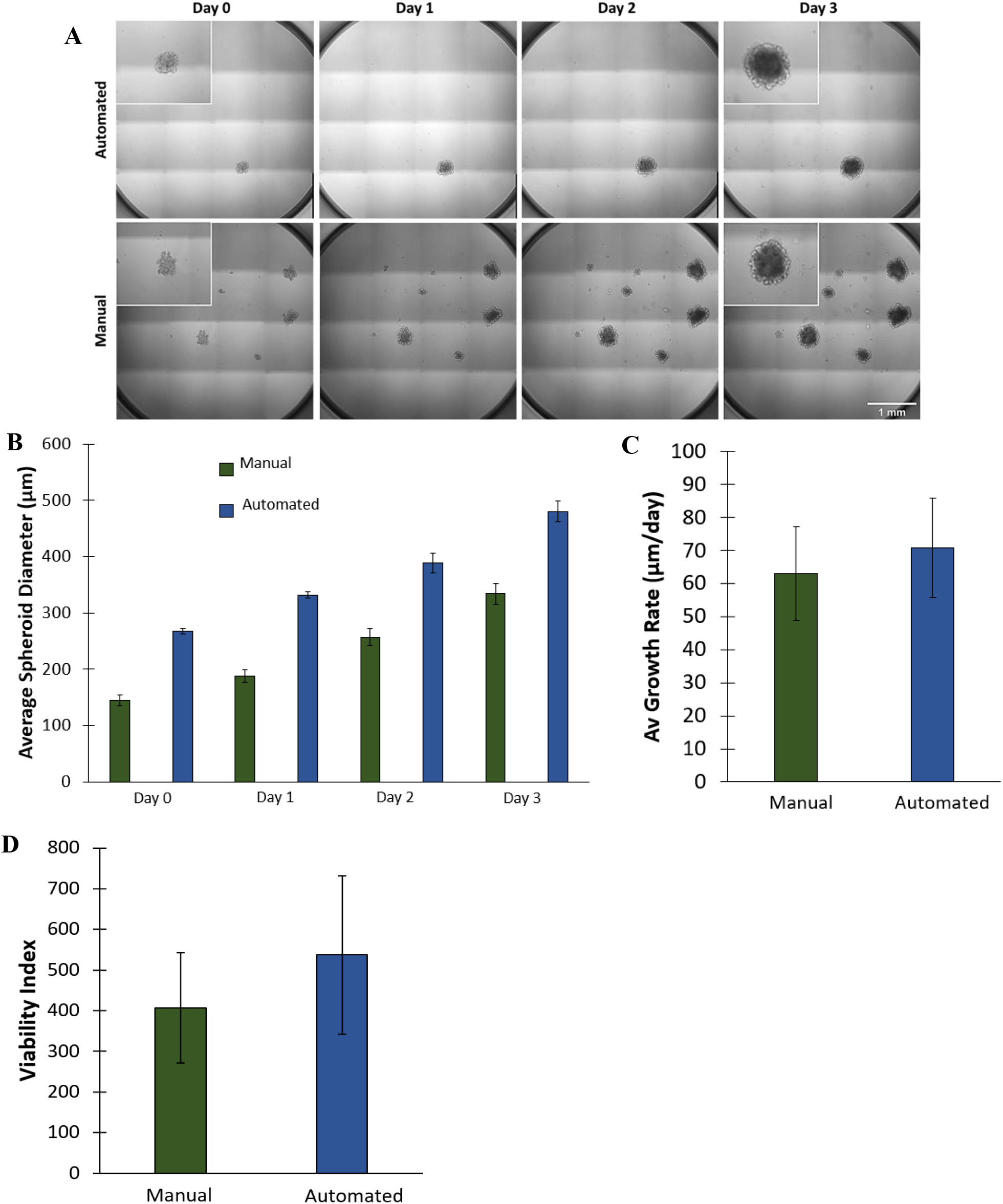
Spheroid morphology. **A)** Transmitted light images of HCT organoids in domes. Insets are enlarged views of select spheroids at day 0 and day 3. **B)** Spheroid size over time in culture. Spheroid diameters from the automated workflow were significantly larger than those from the manual workflow (p ≤ 0.01, Mann-Whitney test). **C)** Growth rate of spheroids seeded via both manual and automated dispensing. Not statistically different. **D)** Spheroid viability after 3 days in culture. Not statistically different (Student’s T test, n = 30 (manual) or 16 (auto), P>0.05).

### Organoid Viability

To assess organoid viability, ATP production (CellTiter-Glo 3D) was measured in spheroids cultured for three days by either the automation system or an experienced user. A viability index was calculated from luminesce values for each well, which were normalized to both spheroid number per well and average spheroid size (diameter) within that well (**Figure 3D**). While there was an apparent higher viability index for the spheroids managed automatically as compared to those managed manually, this difference was not statistically significant.

### PDO Drug Screening Assay

Patient-derived colorectal cancer organoids were treated with a serial dilution of either Adavosertib or Trametinib and the outcomes were compared to no treatment (control). Two different donors were used to assess donor-specific differences in responses. Clear differences could be observed with varying concentrations of drug treatment. Donor 1 PDOs exhibited consistent decreases in ATP production with increasing concentrations of Trametinib and Adavosertib, with the exception of 0.5nM Adavosertib which lowered ATP comparably to higher concentrations (**Figure 4**). Otherwise, low concentrations of each drug were comparable to untreated controls. Donor 2 PDOs exhibited decreased ATP production with all concentrations of Trametinib, but only the highest concentration of Adavosertib resulted in low ATP production (**Figure 4**). Little difference in organoid area was measured between treatment groups of PDOs from either donor (**Figure 5**).

**Figure 4.**
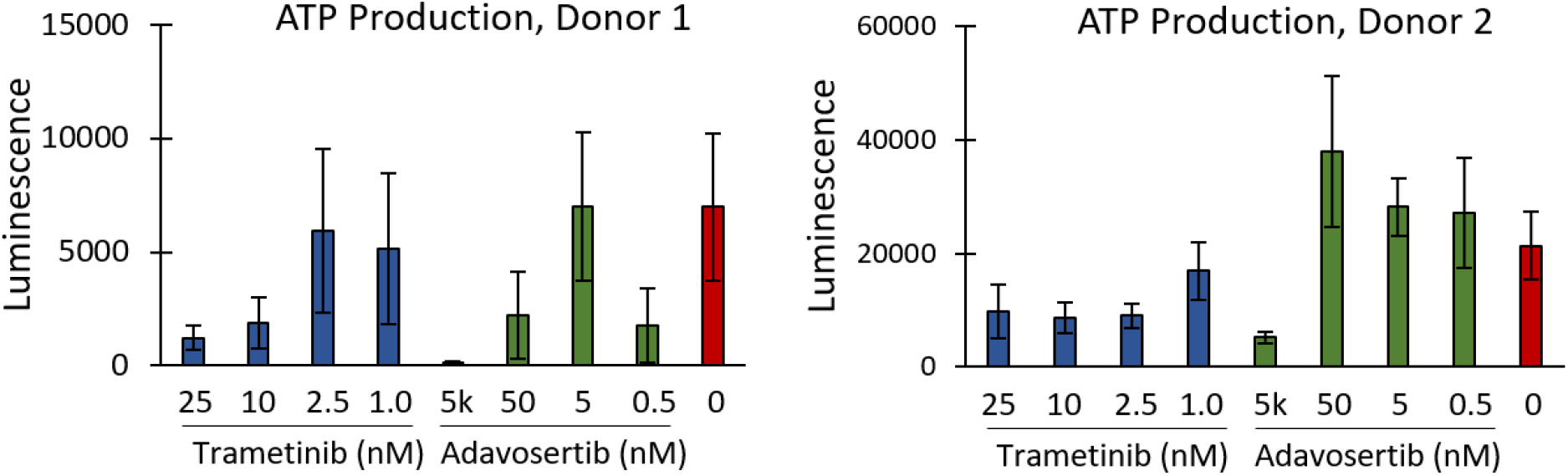
ATP production of PDOs, as measured by CellTiter-Glo® 3D, performed at the end of the assay.

**Figure 5.**
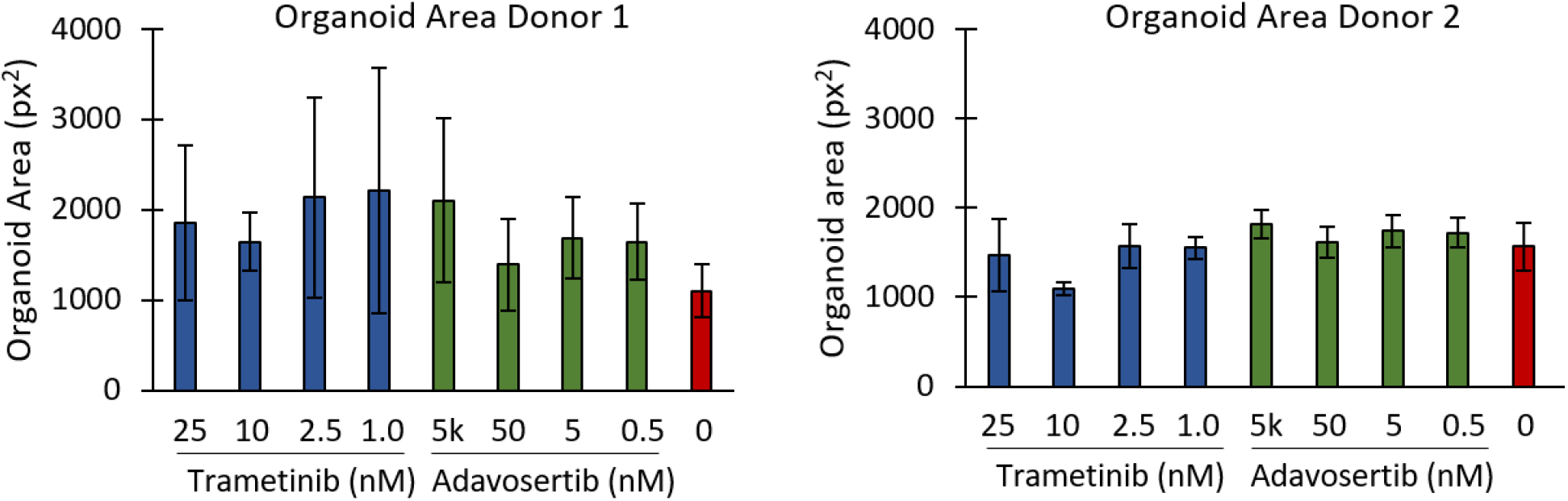
Area of PDOs treated with different concentrations of Trametinib or Adavosertib, compared to untreated controls. Plates were seeded with PDOs from one of two different donors. Bars are mean ± SD

To determine if there were other phenotypic changes in the PDOs in response to the added drugs, we extracted up to 76 features per organoid from the image analysis step and used these for multiparametric analysis. As stated above, little difference was observed in the organoid area across different drug treatments. All phenotypic features representing organoid area, form factor, intensity, and entropy were used to generate principal components. While a variety of analytical options are possible with the StratoMineR platform, we focused on PCA (generalized weighted least squares) followed by Euclidean distance analyses as readouts for this automation demonstration. Principal components extracted from the automatically generated images identified differential responses by the PDOs to the inhibitors (**Figure 6**). PDOs from Donor 1 had a limited response to the two inhibitors (**Figure 6**). In contrast, PDOs from Donor 2 exhibited clear differences in inhibitor responses in a dose-dependent manner (**Figure 6**). An evaluation of hit rate via Euclidean distance score to the controls confirms that Donor 1 PDOs were relatively unresponsive to the 2 inhibitors (hit rate of 31.5%, **Figure 7**), while both inhibitors had a significant impact on Donor 2 PDOs (hit rate of 100%, **Figure 7**).

**Figure 6:**
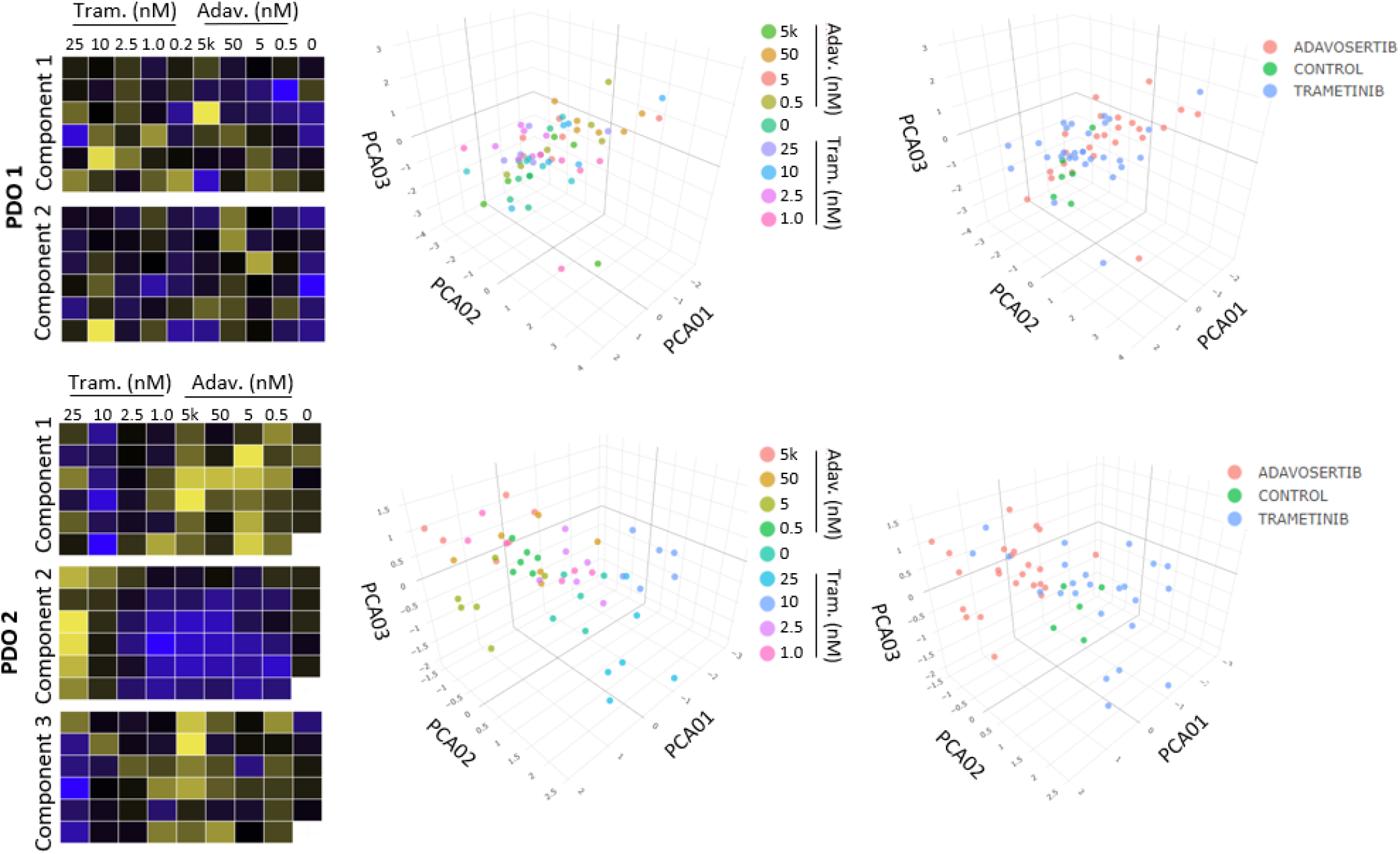
Heat maps and PCA. For Donor 1 (PDO 1), 2 components are shown, which show limited clustering within or between drug concentrations. Donor 2, however, had clear clustering in components 1, 2, and 3, within each drug compared to controls.

**Figure 7.**
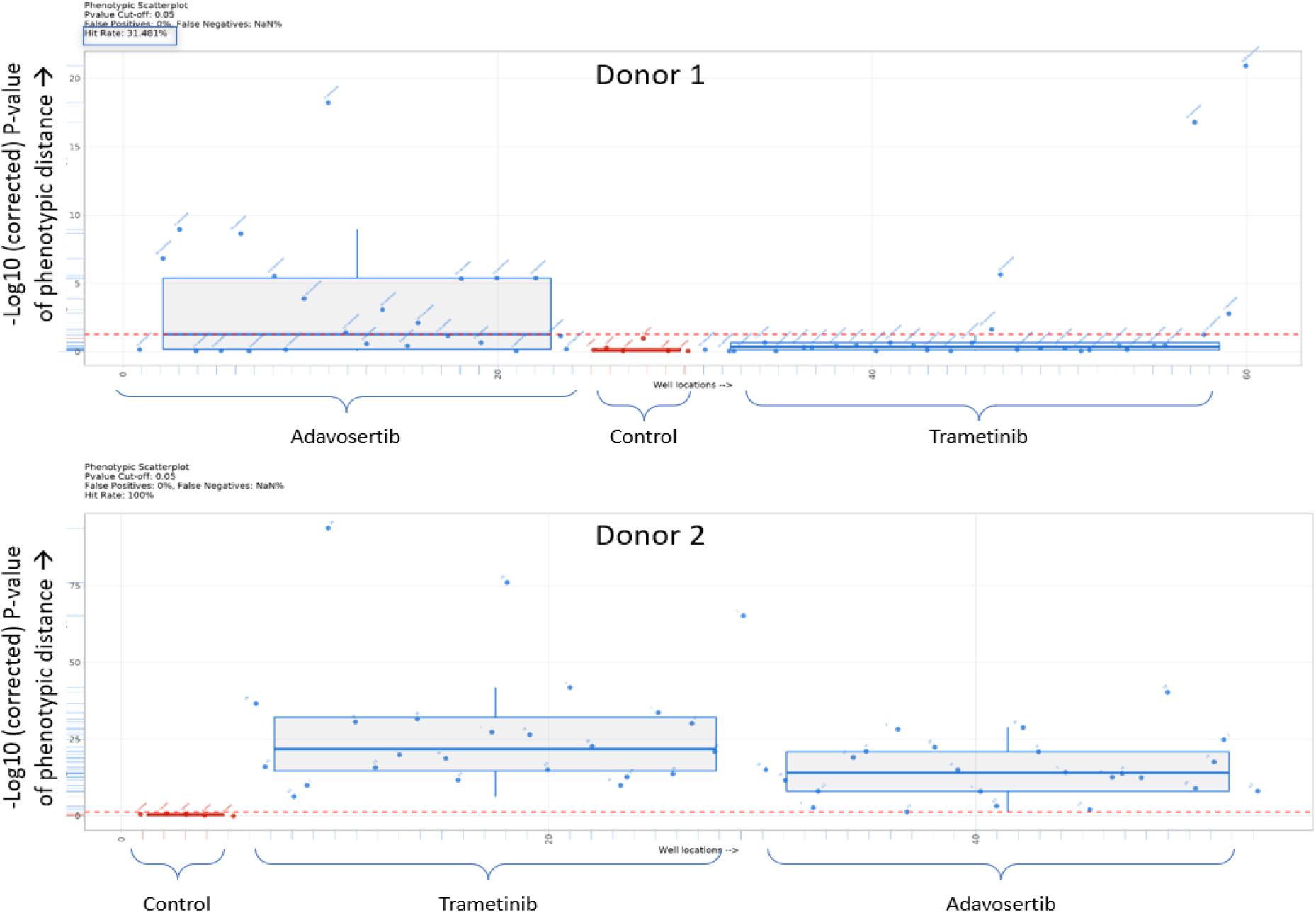
Box plots showing the Euclidean distance from the median of the control (untreated) for PDO 1 and PDO 2. Everything above the red dotted line has a p value <0.05.

## Discussion

Here, we demonstrate an automated workflow in which cultures of colorectal cancer spheroids/organoids are established, maintained, and imaged autonomously via the BioAssemblyBot Platform. The workflow entails all requisite operations, including the formation of Matrigel domes housing the spheroids and interfacing with peripheral instruments. Importantly, spheroid domes were not damaged during the workflow, including with daily media changes, resulting in consistently obtaining expected spheroid sizes and numbers per well.

At the heart of the automation platform is the BioAssemblyBot® 400, a 6-axis robotic arm used routinely in high-precision manufacturing, housed within a framework uniquely suited for the fabrication, manipulation, and manufacturing of living systems, including spheroid/organoid and advanced 3D tissue models. Via a universal interface, the arm can utilize a wide spectrum of end effector tools (BAB Hands), to perform varied tasks and operations, including pipetting and manipulating well plates. This functional flexibility, managed by the user via an intuitive all-in-one software interface called BioApps™, affords the user considerable freedom in performing automated or semi-automated workflows, regardless of complexity. While this current application involved plate management (to multiple operational locations/stations) and precise pipetting, other applications have included 3D bioprinting, cell patterning, and fugitive molding, for example. This allows us to expand our automation platform to a wide range of other tissue fabrication applications beyond the present. Furthermore, while a single plate was worked up for this study, the movement and material transfer capabilities of the robotic arm are readily amenable to conveyor and stacker solutions for continuous spheroid-related operations involving multiple plates, thereby increasing throughput.

Matrigel can be challenging to handle due to its high viscosity and tendency to gel above 4°C. Others have automated Matrigel handling through a variety of methods, although all have slightly different applications and would not be directly suited to dispensing PDO-laden domes. Jiang *et al*. for example, uses a microfluidic system to form individual organoids by combining Matrigel and cells. Then, they use a specialized printer to deposit these Matrigel-based organoids^9^. Reyes-Furrer *et al*. developed a “droplet-on-demand” microvalve system to deposit drops of Matrigel into a dumbbell shape for skeletal muscle modeling^10^. However, neither of these methods are ideal for our application. Here, we required the dome shape because 1) we were pipetting into 96-well plates, which are small, round, wells, 2) a dome shape is more conducive to imaging protocols than filling the well and creating a meniscus, and 3) it uses considerably less material than coating the bottom of the well. To do this, we use the BAB Hand™ | Pipette, which is essentially functioning as a human hand with a pipette, but with the ability to dispense at a more consistent, controllable, rate. Beyond normal pipetting, PDOs in Matrigel can be even more challenging to properly dispense, as this must be done precisely in the center of the well to form a “dome” shape. It takes considerable practice for a scientist to achieve a high success rate of dome formation. In this experiment, use of the BAB improved dome formation rate by 10% (with no practice needed) and dome dispending time was reduced by 50% compared to an experienced scientist. This demonstrates that BAB can improve speed and consequently throughput, without compromising dome formation. In addition to the time savings afforded by the automation, spheroids were not damaged, the expected number and size of spheroids per dome was attained, and there was improved consistency in dome formation.

Trametinib, a MEK1/2 inhibitor, is a well-established anticancer medication. It halts the cell cycle, prevents proliferation, and can induce apoptosis^11, 12^, although its effectiveness can vary based on specific mutations in each individual cancer^13^. Adavosertib (WEE1 inhibitor) similarly has been shown to decrease tumor cell viability in some cancers^14^, although it has not been widely used on colorectal cancers specifically. When treated with these drugs, Donors 1 and 2 had different responses. CellTiter-Glo 3D analysis showed that Donor 1 had an overall lower response to Trametinib, although higher doses were more effective. Donor 2 had a stronger response to Trametinib at both higher and lower doses, shown by reduced viability. Adavosertib was effective only at the highest dose in Donor 2 but was effective across more doses in Donor 1. PCA yielded similar results, but with additional insights. This data indicated that Donor 1 was less responsive to both medications, while Donor 2 responded in a more dose-dependent manner. Combined, this highlights the strong need for the utilization of PDOs in planning patient treatment. Every tumor responds differently to every drug, and there is no one-size-fits-all solution. Combining high-content multiparametric image analysis with PDO screening further strengthens our ability to identify the most effective anti-cancer therapeutics for each particular patient. PCA examines multiple factors together, identifying trends that traditional analysis, which examines one factor at a time (i.e. organoid area, ATP production).

Here, we demonstrate a fully integrated automation platform for screening agent effects on patient-derived tumor organoids. In this use case, the automation platform involved the BioAssemblyBot performing operational tasks in coordination with a modular incubator and high-content imaging systems. Importantly, PDO images were acquired as part of the automation. PDO plates were transferred to the imager and a pre-determined imaging protocol was executed all directed by the PDO BioApps software module. As expected, the automated PDO assay identified patient-specific differences in PDO responses to select signaling inhibitors via high-content image analyses. While a more expanded study is required to establish measurements of consistency, accuracy, and precision, the BAB automation produced results comparable to the work performed by a skilled scientist in approximately half the time with improved efficiencies (e.g. in dome formation). The combination of automated PDO formatting (domes), culturing, treating, and imaging with high-content analysis creates a powerful system for high-throughput drug screening that is highly adaptable yet easy to implement.

## References

1. Moss SM, Schilp J, Yaakov M, et al. Point-of-use, automated fabrication of a 3D human liver model supplemented with human adipose microvessels. SLAS Discov 2022;27:358–368.

2. Liu L, Yu L, Li Z, Li W, Huang W. Patient-derived organoid (PDO) platforms to facilitate clinical decision making. Journal of Translational Medicine 2021;19:40.

3. Fusco P, Parisatto B, Rampazzo E, et al. Patient-derived organoids (PDOs) as a novel in vitro model for neuroblastoma tumours. BMC Cancer 2019;19:970.

4. Aberle MR, Burkhart RA, Tiriac H, et al. Patient-derived organoid models help define personalized management of gastrointestinal cancer. Br J Surg 2018;105:e48–e60.

5. Lee SH, Hu W, Matulay JT, et al. Tumor Evolution and Drug Response in Patient-Derived Organoid Models of Bladder Cancer. Cell 2018;173:515–528 e517.

6. Pasch CA, Favreau PF, Yueh AE, et al. Patient-Derived Cancer Organoid Cultures to Predict Sensitivity to Chemotherapy and Radiation. Clin Cancer Res 2019;25:5376–5387.

7. Caicedo JC, Cooper S, Heigwer F, et al. Data-analysis strategies for image-based cell profiling. Nature Methods 2017;14:849–863.

8. Omta WA, van Heesbeen RG, Pagliero RJ, et al. HC StratoMineR: A Web-Based Tool for the Rapid Analysis of High-Content Datasets. Assay Drug Dev Technol 2016;14:439–452.

9. Jiang S, Zhao H, Zhang W, et al. An Automated Organoid Platform with Inter-organoid Homogeneity and Inter-patient Heterogeneity. Cell Rep Med 2020;1:100161.

10. Alave Reyes-Furrer A, De Andrade S, Bachmann D, et al. Matrigel 3D bioprinting of contractile human skeletal muscle models recapitulating exercise and pharmacological responses. Commun Biol 2021;4:1183.

11. Elbadawy M, Sato Y, Mori T, et al. Anti-tumor effect of trametinib in bladder cancer organoid and the underlying mechanism. Cancer Biol Ther 2021;22:357–371.

12. Zeiser R, Andrlova H, Meiss F. Trametinib (GSK1120212). Recent Results Cancer Res 2018;211:91–100.

13. Hirashita Y, Tsukamoto Y, Kudo Y, et al. Early response in phosphorylation of ribosomal protein S6 is associated with sensitivity to trametinib in colorectal cancer cells. Laboratory Investigation 2021;101:1036–1047.

14. Kopper O, de Witte CJ, Lohmussaar K, et al. An organoid platform for ovarian cancer captures intra- and interpatient heterogeneity. Nature medicine 2019;25:838–849.

